# Impact of ionizing radiation on the environmental microbiomes of Chernobyl wetlands

**DOI:** 10.1101/2022.01.17.476627

**Authors:** Elin Videvall, Pablo Burraco, Germán Orizaola

**Author notes:** Correspondence Germán Orizaola, Zoology Unit, Department of Biology of Organisms and Systems, University of Oviedo, c/ Catedrático Rodrigo Uría s/n, 33071 Oviedo-Asturias, Spain. Elin Videvall and Pablo Burraco contributed equally to the study.

## Abstract

Radioactive contamination in the form of ionizing radiation can be a devastating pollutant because it has the potential to cause damage to DNA and other biomolecules. Anthropogenic sources of ionizing radiation include accidents in nuclear power plants, such as the one in Chernobyl 1986, which caused long-term radioactive pollution. Studies on animals within radioactive zones have provided us with a greater understanding of how wildlife can persevere despite chronic radiation exposure, however, we still know very little about the effects of radiation on the microbial communities in the environment. Here, we examined the impact of ionizing radiation and other environmental factors on the diversity and composition of environmental microbiomes in the wetlands of Chernobyl. We combined extensive field sampling along a gradient of radiation together with 16S rRNA high-throughput metabarcoding (Illumina NovaSeq). While radiation did not affect the alpha diversity of the microbiomes in sediment, soil, or water, it had a strong effect on the beta diversity, indicating that the microbial composition was affected by ionizing radiation. Specifically, we detected several microbial taxa that were more abundant in areas with high radiation levels within the Chernobyl Exclusion Zone, including bacteria and archaea known to be radioresistant. Overall, our results reveal the existence of rich and diverse microbiomes in Chernobyl wetlands, with multiple taxonomic groups that are able to thrive despite the radioactive contamination. Further field and laboratory-based approaches will help to forecast the functionality and re-naturalization dynamics of radiocontaminated environments.

## 1 INTRODUCTION

Human activities are transforming natural ecosystems at an unprecedented rate (Steffen et al., 2007). The intense use and transformation of natural habitats over the last decades have had a severe impact on biodiversity (Palumbi, 2001). Habitat destruction, climate alteration, invasive species, and the release of numerous pollutants into the environment are the main factors behind biodiversity decline (Rands et al., 2010). Pollutants, in particular, can affect species distribution and abundance, lead to the extirpation of the most susceptible ones, and alter biological functions, ecological networks and ecosystem services (Edwards, 2002).

Ionizing radiation is a rare but potentially devastating pollutant. This type of radiation is present in the environment at low levels as a natural phenomenon (e.g. cosmic and terrestrial radiation), and generally does not cause damage to living organisms. However, certain human activities, such as weapons testing and accidents at nuclear power plants, can involve releases of ionizing radiation above safety levels. Ionizing radiation may damage organic molecules, including DNA, and cause malfunctions in cell processes that lead to cellular and organismal death (Han & Yu, 2010; Reisz et al., 2014). Indeed, the effects of exposure to acute levels of ionizing radiation are acknowledged to negatively impact organisms and ecosystems (Møller & Mousseau, 2006, 2015).

The accident at the Chernobyl nuclear power plant, on the 26^th^ of April 1986, led to the largest release of radioactive material in human history (UNSCEAR, 1988). An exclusion zone of ca. 4,700km^2^ was created around the power plant (Chernobyl Exclusion Zone, CEZ), preventing human settlement in the area, conditions that remains in effect. Exposure to the acute radiation levels generated by the Chernobyl accident caused a severe impact on the organisms in the area, including humans (Smith & Beresford, 2005; Møller & Mousseau, 2015). Studies on wildlife conducted after the accident reported that the radioactive contamination led to reductions in species diversity, multiple physiological costs, and increased DNA damage (Møller & Mousseau, 2006, 2015). However, the effects of ionizing radiation are far from general; while some studies reported negative consequences on wildlife populations currently living in the area (e.g. Beaugelin-Seiller et al., 2020), others have reported population recoveries (e.g. Deryabina et al., 2015; Schlichting et al., 2019), and signs of adaptation to the chronic exposure (e.g. Galvan et al., 2014; Møller & Mousseau, 2016). There is still an intense scientific debate about the long-lasting effects of chronic exposure to moderate levels of ionizing radiation on biodiversity (e.g. Møller & Mousseau, 2006; Beresford et al., 2016, 2020).

Bacterial communities are crucial for maintaining ecosystem functions due to their role in the cycling, retention, and release of major nutrients and soil carbon (Gucht et al., 2007; Newton et al., 2011; McKenney et al., 2018). Chronic exposure to pollutants, including ionizing radiation, can compromise the diversity and composition of bacterial communities (Chapin et al., 2000; Ager et al., 2010). A host-associated microbiome is predominantly constrained by the bacteria they are able to recruit from their environment, and the composition and diversity of the resulting microbial community in the host can have important effects on their health (Liu et al., 2019). Changes in the composition of environmental microbiomes as a consequence of chronic exposure to radiation can therefore have indirect effects on local wildlife by changing their capacity to acquire specific symbionts from the environment.

Despite the crucial ecological role of microbes in the environment, the impact of ionizing radiation on environmental microbiomes has not been comprehensively explored, and the Chernobyl accident represents an ideal opportunity in this regard (IAEA, 2006). Bacteria are often considered to have a greater resistance to ionizing radiation than other organisms (ICRP, 2014). Some bacterial taxa have been recovered from highly radio-contaminated environments, and, in some cases, their radioresistance capacity has been demonstrated under laboratory conditions (Ryabova et al., 2020). However, these studies have been restricted to a handful of taxa while the majority of environmental bacteria have never been studied in relation to radiation, neither in the laboratory nor in their natural environment. Shortly after the Chernobyl accident, studies on soil samples from the central area of the Chernobyl Exclusion Zone reported a two-fold lower abundance in bacteria, compared to control non-contaminated areas outside the Zone (Romanovskaya et al., 1998; Yablokov, 2009). Some soil bacteria from the Chernobyl area were able to accumulate high doses of radioactive substances (e.g. ^137^Cs), as in the case of Agrobacterium sp., Enterobacter sp., and Klebsiella sp. (Yablokov, 2009). Chapon et al. analysed highly contaminated areas in the CEZ and identified a high diversity of soil bacteria using a mix of culturing techniques and sequencing tools (Chapon et al., 2012), whereas Hoyos-Hernandez et al. identified genes potentially associated with radiation resistance in prokaryotes (Hoyos-Hernandez et al., 2019). Recent studies have also examined the effects of radiation on the gut microbiome of wild vertebrates (e. g. Ruiz-González et al., 2016, Lavrinienko et al., 2018a,b) and earthworms (Newbold et al., 2019). Therefore, despite some progress, information on the bacterial communities present along the gradient of radioactive contamination in Chernobyl is still scarce. Acquiring this knowledge is not only relevant for evaluating the impact of radiation on microbes themselves, but also for further comprehensive assessments of the impact of radiation on multicellular organisms’ host-associated microbiota.

In this study, we examined environmental factors affecting the diversity and composition of bacterial communities in wetlands, by analyzing pond water, sediment, and soil at multiple locations inside and outside the Chernobyl Exclusion Zone. We paid particular attention to examining the impact of radioactive contamination on these environmental microbiomes, but also evaluated other variables that might induce changes in bacterial composition. Wetlands cover between 5-10% of the earth’s land surface (Mitsch & Gosselink, 2007) and provide essential ecosystem services such as carbon reservoirs (Mitsch et al., 2013). As in other natural areas, Chernobyl wetlands are essential for a large number of aquatic and terrestrial plants and animals. Understanding how ionizing radiation alters the bacterial communities associated with these environments is crucial for a comprehensive evaluation of radio-contaminated ecosystems.

## 2 MATERIALS AND METHODS

### 2.1 Field work

Sampling was conducted in Northern Ukraine, inside and outside the Chernobyl Exclusion Zone between the 29^th^ of May and the 3rd of June 2019 (Figure 1, Table S1). In total, we selected 21 permanent wetlands: 16 within the Chernobyl Exclusion Zone and 5 in a nearby control area with background radiation levels (Figure 1, Table S1). All the sampled wetlands shared similar characteristics: small to medium size wetlands with reed beds, situated within a matrix of forest and meadows on sandy soils (soddy-podzolized sandy and clay-sandy soils; Soil Map of Ukraine accessed from https://esdac.jrc.ec.europa.eu/content/title-russia-soil-map-ukraine). To examine the diversity and composition of bacterial communities, at each site we collected samples from three environment types: water, pond sediment, and soil near the banks of the wetland. Each *environment type* was sampled at three points within each site. In total, we collected 189 samples using tubed sterile Dryswab MW100 swabs with rayon tip. We collected water samples by swirling a swab on the water surface for 25 seconds in areas about one-meter depth, and about two meters from the shore. Sediment was collected at ca. 0.5 m depth with a sampler that removes the top 10 cm of sediment, and swabs were inserted into the sediment five times for five seconds each time. Soil samples were collected on land, 5-10 meters from the water edge, by first removing the top 5 cm of soil and then twirling a swab inside the exposed soil for 15 seconds. The swabs were placed in individual plastic vials on-site and stored in a portable cooler until arrival at our laboratory in Chernobyl, where samples were stored in a fridge at 4 °C. Finally, we transported the samples to our laboratory at the University of Oviedo (Spain), where they were stored at −20 °C until further processing.

**FIGURE 1.**
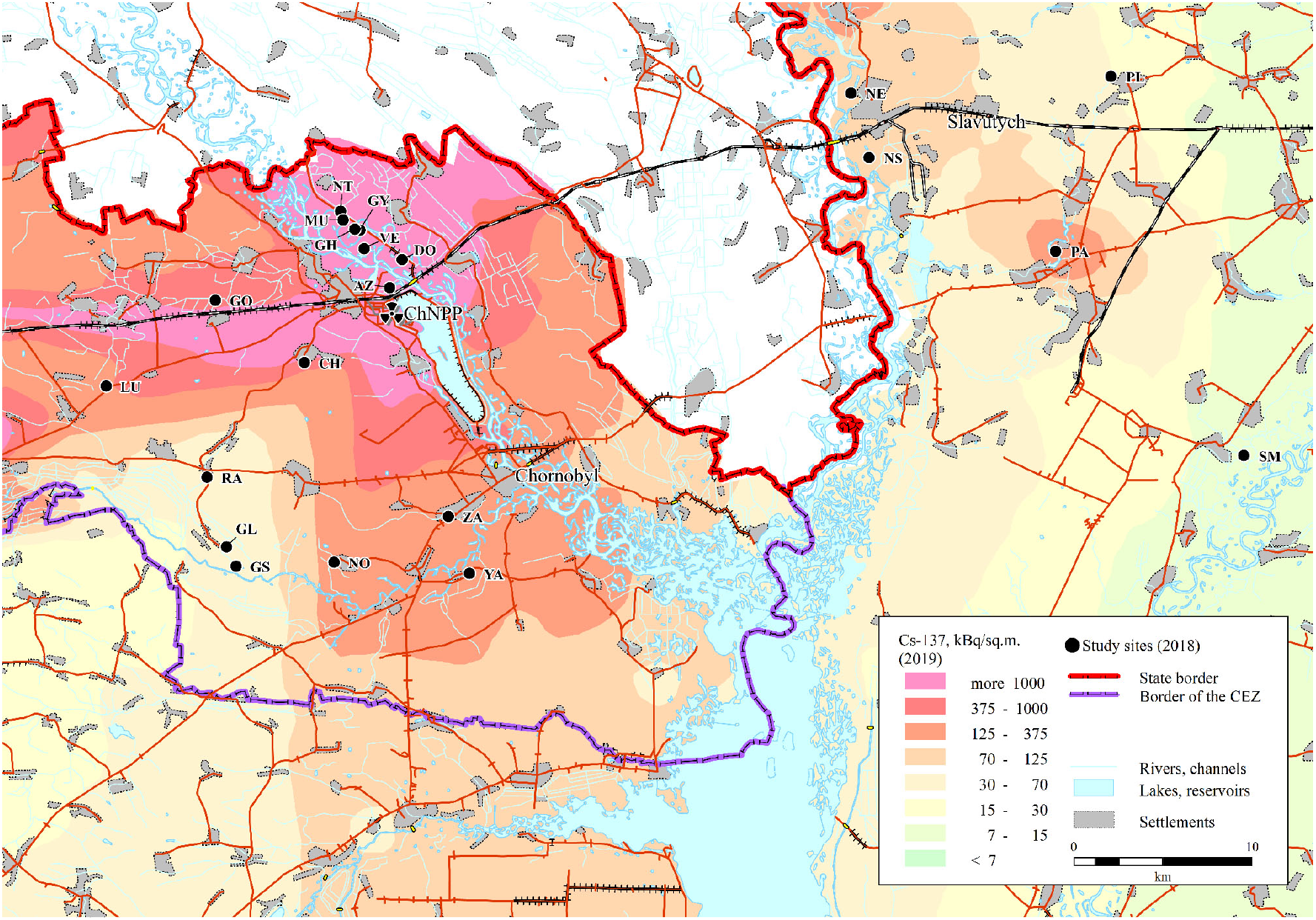
Map showing the sampling sites in Northern Ukraine. Abbreviations refer to the location name (see Table S1 for details). The underlying ^137^Cs soil data (decay corrected to spring 2018) are derived from the Atlas of Radioactive Contamination of Ukraine (Intelligence Systems GEO, 2011).

At each site, we measured water chemical characteristics with a Hanna multiparametric portable meter HI9811-5: temperature, pH, total dissolved solids (TDS; ppm/mg/L), and electrical conductivity (EC; μS/cm), at the same three sampling points where we collect the water and sediment samples. We measured radiation levels (in μSv/h) with an MKS-AT6130 handheld radiometer, at five points next to each environment type and site, by placing the radiometer 5 cm above the sampled area (see Burraco et al., 2021 for details).

### 2.2 DNA isolation, library preparation, and sequencing

We used the DNeasy PowerSoil DNA isolation kit (Qiagen) to isolate DNA from soil and sediment samples, and the NZY Tissue gDNA isolation kit (NZYTech) to isolate DNA from water samples. We resuspended DNA in a final volume of 100 or 50 μL when Qiagen or NZY kit were used, respectively. We included an extraction blank in every DNA extraction round and treated it as a regular sample to check for cross-contamination.

For library preparation, we amplified a fragment of the bacterial 16S rRNA region (V4) of around 300 bp using the standard primers 515F (5’ GTG YCA GCM GCC GCG GTA A 3’) (Parada et al., 2016) and 806R (5’ GGA CTA CNV GGG TWT CTA AT 3’) (Apprill et al., 2015). We ran PCRs using a final volume of 25 μL, containing 2.5 μL of template DNA (except for 11 samples with low concentration for which we used 5 μL), 0.5 μM of the primers, 12.5 μL of Supreme NZYTaq 2x Green Master Mix (NZYTech), and ultrapure water up to 25 μL. The reaction mixture was incubated as follows: an initial denaturation at 95 °C for 5 min, followed by 25 cycles of 95 °C for 30 s, 46 °C for 45 s, 72 °C for 45 s, and a final extension step at 72 °C for 7 min.

For each sampled site, we pooled the three PCR replicates from each environment type (e.g. 3 × water samples at each location). Once pooled, we attached the oligonucleotide indices required for multiplexing in a second PCR with identical conditions as previously but during 5 cycles and using 60 °C as the annealing temperature (see Vierna et al., 2017). We included a negative control in every PCR run to check for contamination during library preparation. The libraries were run on 2% agarose gels stained with GreenSafe (NZYTech), and imaged under UV light to verify the library size. Libraries were purified using the Mag-Bind RXNPure Plus magnetic beads (Omega Biotek), and then pooled in equimolar amounts according to the quantification data provided by a Qubit dsDNA HS Assay (Thermo Fisher Scientific). The pool was sequenced in a fraction of an Illumina NovaSeq paired end 250bp run.

### 2.3 Data processing

We evaluated the quality of the reads using FastQC (Andrews, 2010) in combination with MultiQC (Ewels et al., 2016). We next used DADA2 (Callahan et al., 2016) implemented in QIIME2 (release 2020.2; Bolyen et al., 2018) to remove PCR primers, quality-filter reads, denoise, merge the pairs, remove chimaeric reads, and cluster the resulting sequences into amplicon sequence variants (ASVs). Forward and reverse reads were truncated at position 249 before merging with a minimum overlapping region of 12 identical base pairs. Taxonomy was assigned using a trained classifier of the SILVA reference database (Quast et al., 2013, release 138 December 2019), with the feature-classifier classify-sklearn approach, implemented in QIIME2 (Bokulich et al., 2018). A phylogenetic tree was constructed in QIIME2 (v. 2020.2), using MAFFT (Katoh & Standley, 2013) and FastTree2 (Price et al., 2010) for phylogenetic analyses.

The ASV table was imported into R (v. 4.0.2; R-Team-Core, 2020), and the packages *phyloseq* (v. 1.32.0; McMurdie & Holmes, 2013) and *vegan* (v. 2.5-6; Oksanen et al., 2019) were used for statistical analysis. From downstream analyses, we excluded: the ASVs with a single read in the whole dataset (singletons), the completely unassigned sequences (no bacterial classification), and those from chloroplast and mitochondrial origin. We also removed ASVs occurring at a frequency below 0.01% in each sample to account for potential misassignments during library preparation and low-frequency contaminants. Furthermore, we used decontam (v. 1.6.0; Davis et al., 2018) to identify and eliminate 9 potential contaminating ASVs in the reagents by analyzing the blank samples that were sequenced simultaneously as negative controls. A high and even read depth of high-quality sequences (average number of reads per sample = 41,210) allowed us to safely rarefy the data to 30,000 reads without losing any samples or reducing statistical power.

### 2.4 Statistical Analysis

Sampling sites were assigned to three different areas regarding their location and radiation levels: *CEZ-high* for sites located inside the Chernobyl Exclusion Zone in environments with soil and sediment radiation levels > 2 μSv/h; *CEZ-low* for sites located inside Chernobyl with radiation levels < 0.5 μSv/h; and *Outside-CEZ* for sites outside Chernobyl, where radiation levels were < 0.2 μSv/h (Table S1). We calculated Alpha diversity for the bacterial communities in water, pond sediment, and soil using three different metrics: Richness (i.e. total number of unique ASVs), Shannon index (which takes into account both richness and evenness), and Faith’s index (i.e. phylogenetic diversity). Beta diversity was measured with the Bray-Curtis distance metric, which accounts for both the presence/absence and abundance of microbes, and with unweighted UniFrac, which measures phylogenetic distances between bacteria. We used *betadisper* and *adonis* in the R package *vegan* to calculate homogeneity of group dispersion and to perform PERMANOVAs to assess variation. Radiation levels (μSv/h), Total Dissolved Solids (TDS; ppm/mg/L), and Electrical Conductivity (EC; μS/cm) were log-transformed prior to analysis. Finally, we used ANCOM with bias correction (ANCOM-BC, v. 0.99.1; Lin & Peddada, 2020) on each environment type (water, sediment, soil), to identify specific taxa that were associated with higher or lower radiation levels. In ANCOM-BC, we used a conservative variance estimate and accounted for site location within or outside the CEZ (group parameter).

## 3 RESULTS

We identified a total of 20,816 unique ASVs across the three environment types (surface water, pond sediment, and soil) from the 21 sampled sites. Soil and pond sediment had the highest numbers of ASVs (soil = 11,553, sediment = 11,033), whereas the water samples had much fewer ASVs (water = 3,091; Fig. 2a). Despite the large diversity of microbes in both soil and pond sediment, the vast majority of ASVs was not shared across the sample types (Figure 2a). Water had substantially lower bacterial diversity (25% lower) and richness (70% lower) than sediment and soil (*P* < 0.001, in all cases). There were no differences in alpha diversity among the sites located within the CEZ (high and low radioactivity) and outside the CEZ, for any environment type (sediment, soil, water) (ANOVA of Shannon index: sediment, *F*_2,16_ = 0.29, *P* = 0.755; soil, *F*_2,16_ = 1.51, *P* = 0.252; water, *F*_2,16_ = 0.55, *P* = 0.588; Fig. 2b). Similar results were obtained for ASV richness (sediment, *F*_2,16_ = 0.12, *P* = 0.891; soil, *F*_2,16_ = 1.24, *P* = 0.315; water, *F*_2,16_ = 0.37, *P* = 0.698; Fig. 2c) and Faith’s phylogenetic diversity (sediment, *F*_2,16_ = 0.18, *P* = 0.834; soil, *F*_2,16_ = 0.41, *P* = 0.668; water, *F*_2,16_ = 0.17, *P* = 0.847; Fig. 2d). The effects of temperature and pH were not significant for any of these analyses (*P* > 0.159, in all cases). We also found no correlation between radiation levels and bacterial diversity and richness in either of the three sample types (*P* > 0.32, in all cases; Figure S1).

**FIGURE 2.**
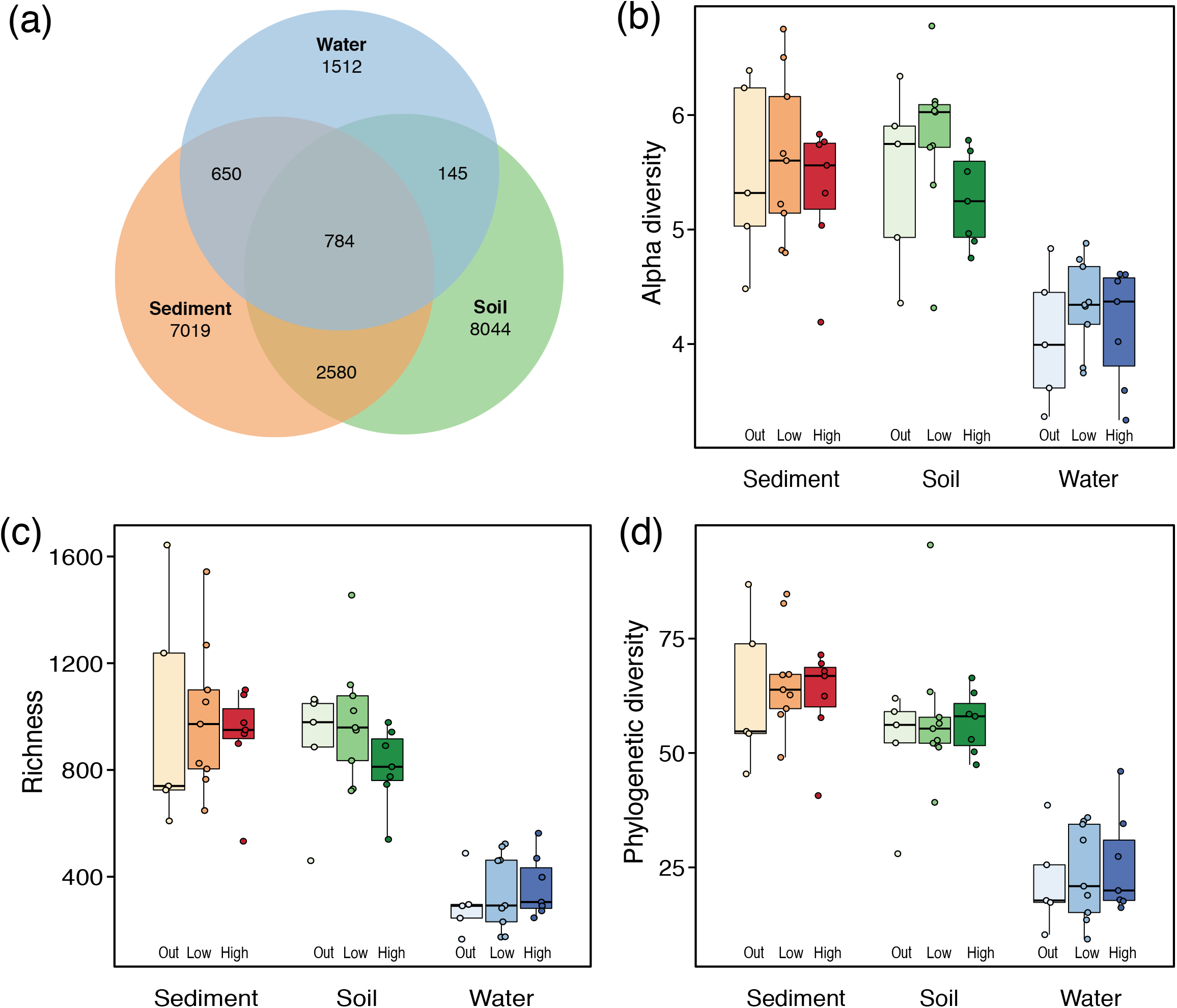
Composition of bacterial communities in wetlands within and outside Chernobyl. (a) Number of unique and shared amplicon sequence variants (ASVs) in each environment type, (b) alpha diversity, (c) richness, and (d) phylogenetic diversity.

The composition of environmental microbiomes differed substantially between the three environments (*R*^*2*^ = 17.5%, *P* < 0.001), with the water microbiome being the most differentiated (Figure 3a). Analyses of sources of variance for each of the sample types using Bray-Curtis distances showed that radiation influenced the composition of both the sediment microbiome (*R*^*2*^ = 6.8%; Table 1, Figure 3b) and the soil microbiome (*R*^*2*^ = 7.5%; Figure 3c). Temperature significantly affected both the water microbiome (*R*^*2*^ = 8.2%) and the soil microbiome (*R*^*2*^ = 7.7%; Table 1), whereas dissolved solids had an effect on the water microbiome composition (*R*^*2*^ = 10.3%; Table 1). While radiation levels explained a decent proportion of the water microbiome variance (*R*^*2*^ = 6.3%; Figure 3d), this effect was non-significant (*P* = 0.066). In contrast, when analyzing microbiome composition with a phylogenetic distance metric (UniFrac), radiation showed strong significant effects on both soil (*R*^*2*^ = 7.2%, *P* = 0.004) and water samples (*R*^*2*^ = 7.8%; *P* = 0.009), but not sediment microbiome (*R*^*2*^ = 6.3%, *P* = 0.08; Table S2). All group dispersion tests (function *betadisper* within vegan package) discarded any significant heterogeneity of environment microbiomes (*P* > 0.1), indicating that the beta diversity differences in Table 1 and Figure 3 could not be explained simply by sample dispersion.

**FIGURE 3.**
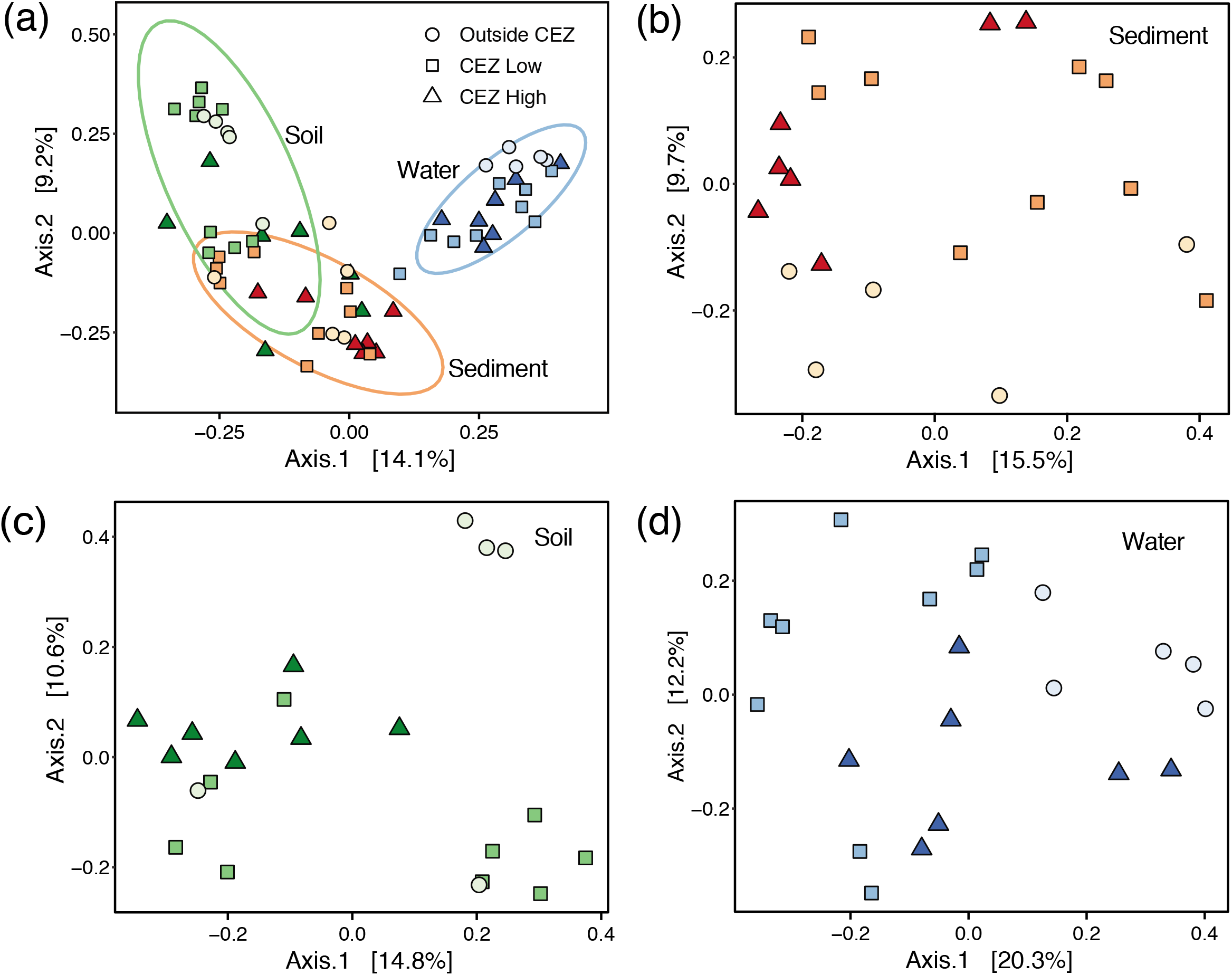
Principal Coordinate Analysis (PCoA) on Bray–Curtis dissimilarity distances of the microbiomes in (a) all samples colored by environment type; and each environment type: (b) sediment, (c) soil, and (d) water.

**TABLE 1.** Permanova on the effects of radiation and environmental factors of the three environment microbiomes using Bray-Curtis distances. TDS = Total Dissolved Solids, EC = Electrical Conductivity.

|  | Sediment |  |  | Soil |  |  | Water |  |  |
| --- | --- | --- | --- | --- | --- | --- | --- | --- | --- |
|  | *F* | *R*<sup>2</sup> | *P* | *F* | *R*<sup>2</sup> | *P* | *F* | *R*<sup>2</sup> | *P* |
| Radiation | 1.420 | 0.068 | 0.035 | 1.567 | 0.075 | 0.013 | 1.466 | 0.063 | 0.066 |
| Temperature | 1.049 | 0.050 | 0.377 | 1.599 | 0.077 | 0.016 | 1.910 | 0.082 | 0.008 |
| pH | 1.245 | 0.060 | 0.133 | 0.950 | 0.046 | 0.562 | 1.603 | 0.069 | 0.055 |
| TDS | 1.222 | 0.059 | 0.132 | 0.834 | 0.040 | 0.785 | 2.415 | 0.103 | 0.002 |
| EC | 0.921 | 0.044 | 0.651 | 0.859 | 0.041 | 0.762 | 0.968 | 0.041 | 0.468 |

The taxonomic analysis of environment microbiomes showed that sediment and soil communities had a diverse composition consisting of multiple abundant phyla, with Actinobacteriota more abundant in soil and Desulfobacterota more common in sediment (Figure 4). Water communities consisted almost exclusively of Proteobacteria and Bacteroidota (Figure 4). The effects of radiation on specific members of the microbiomes evaluated using ANCOM-BC revealed several bacterial families with higher abundances in the localities with higher radioactivity (Figure 5). The majority of the taxa that were positively associated with radiation levels were unique to their respective environment microbiome; for example, *Prolixibacteraceae* in soil, *Methylococcaceae* in sediment, and *Rhodocyclaceae* in water. However, some taxa were differentially abundant across sample types, e.g. *Lentimicrobiaceae* was more abundant at the high-radiation localities in both soil and water microbiomes, and *Eubacterium coprostanoligenes* group was more abundant at the high-radiation localities in both sediment and water. Families associated with higher abundance in high-radiation areas also included *Anaerolineae* and Thermoplasmata in sediment or *Smithellaceae* and *Geobacteraceae* in water; whereas families with lower abundances in high-radiation soil microbiomes included *Micromonosporaceae*, *Microbacteriaceae*, TK10, *Fibrobacteraceae*, and *Chthoniobacteraceae* (Figure 5).

**FIGURE 4.**
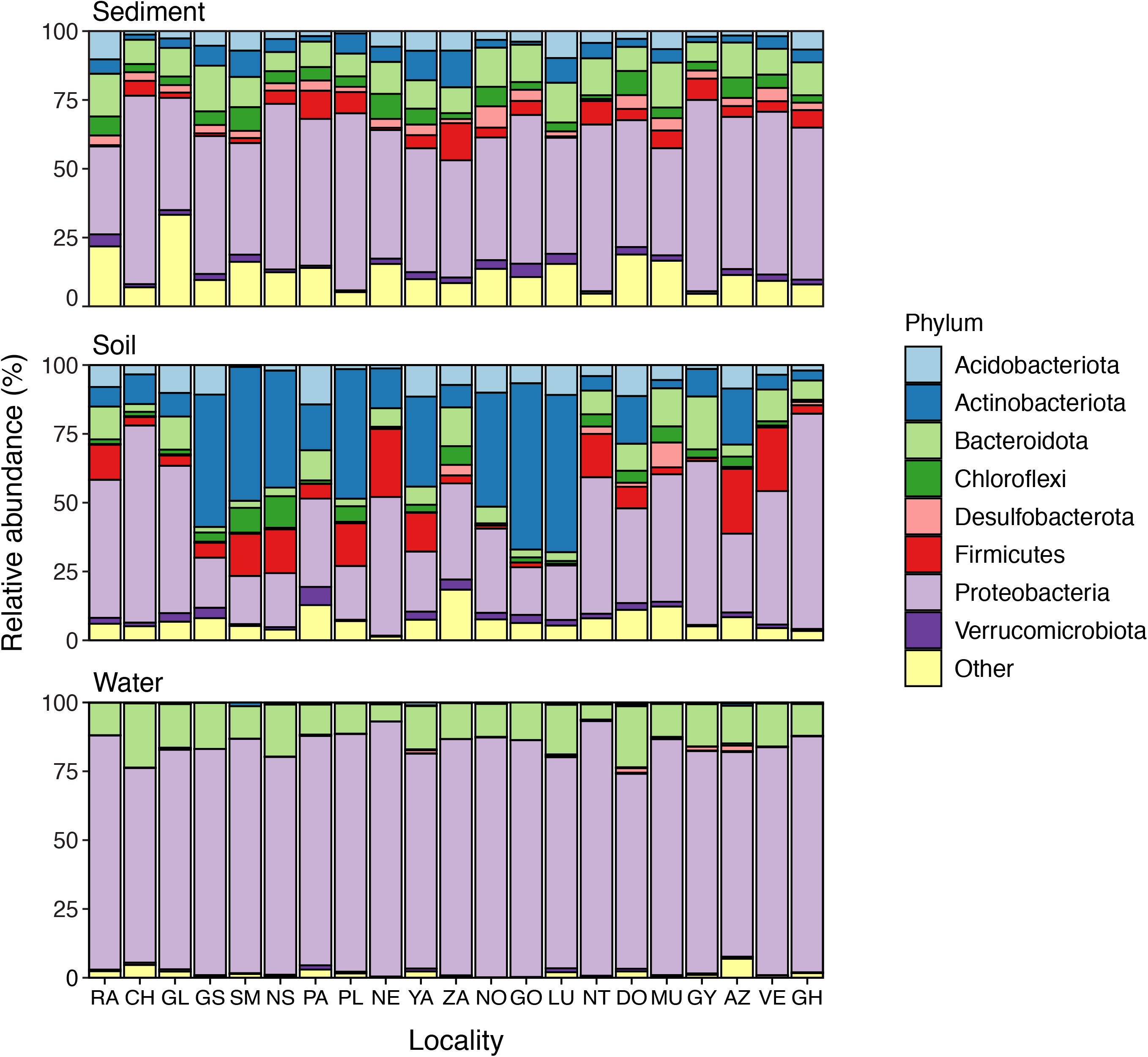
Taxonomic composition of Chernobyl wetland microbiomes in sediment, soil, and water. Localities (x-axis) are arranged according to radiation levels with the right-most localities having the highest radioactivity.

**FIGURE 5.**
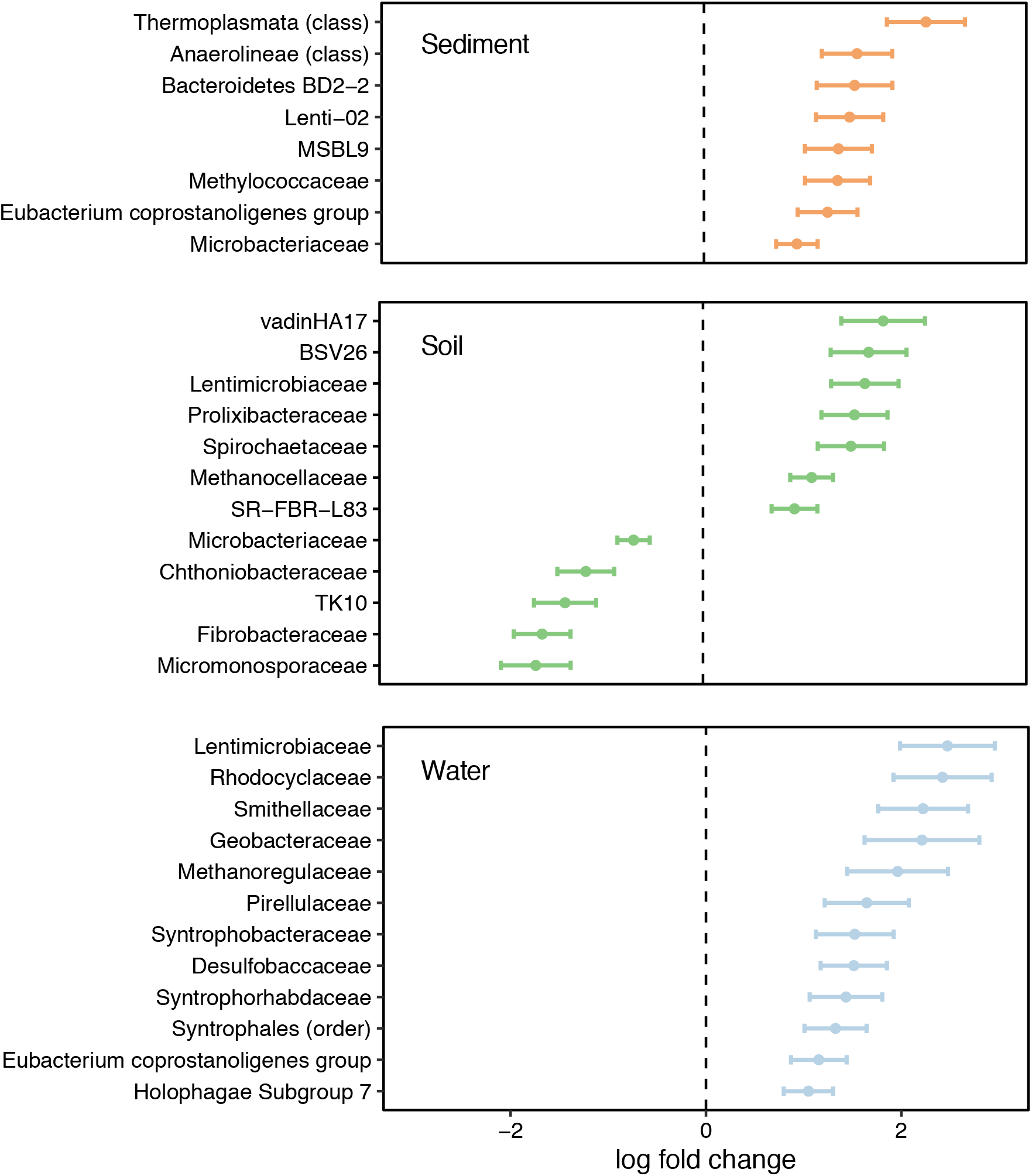
Differentially abundant taxa in Chernobyl wetland microbiomes in response to radiation levels in sediment, soil, and water. The error bars show the unstandardized effect size (beta) ± standard error (SE). Y-axis lists significantly differentially abundant bacterial families (or the closest taxonomic order assigned). Positive log fold change indicates higher abundance in localities with higher radioactivity.

## 4 DISCUSSION

Chernobyl wetlands maintain rich and diverse microbial communities three decades after the accident in the nuclear power plant. Our study reveals that the diversity and richness of bacterial communities in pond sediment, shore soil, and water were similar between wetlands sampled inside and outside the Chernobyl Exclusion Zone, and that these two parameters were not influenced by radiation levels. However, the beta diversity and composition of the bacterial communities in Chernobyl wetlands were affected by different environmental factors, with radioactivity being the main effect. In addition, we discovered several bacterial families with higher abundance in sites with high radiation levels. Understanding how bacterial communities are influenced by radioactive contamination is crucial for evaluating the impact of the Chernobyl accident in the environment and to forecast the future development of the ecosystems in the area.

Our study presents the most comprehensive assessment of bacterial communities in areas affected by the Chernobyl accident to date, reporting more than 20,000 unique ASVs. It effectively builds upon previous studies of the area, which used other techniques rather than metabarcoding to identify bacteria (Theodorakopoulos et al., 2017; Lavrinenko et al., 2018a,b; Chapon et al., 2012). By using metabarcoding, we found substantial differences between environments, with very high bacterial richness and diversity across all soil and sediment microbiomes. These results corroborate the findings in the Earth Microbiome Project (EMP), where sediment and soil microbiomes were shown to vastly outnumber all other free-living microbial communities in terms of bacterial richness (Thompson et al., 2017). Similar to the EMP, our water microbiomes had much lower richness and differed in community composition. Stark microbial differences across environment types are not surprising given their highly differentiated characteristics (e.g. microclimate and environmental components).

Bacterial diversity and richness of the studied sites did not change across the gradient of radioactive contamination, or between samples collected inside Chernobyl and in areas with background radiation levels outside the Exclusion Zone. This finding agrees with previous studies that sampled biofilm communities and bacteria in trenches for radioactive waste disposal in Chernobyl, both reporting that the diversity of bacteria did not change between sites with high and low levels of radiation, or even when compared with remote, non-contaminated areas, although these studies were only able to examine a low number of OTUs (Ragon et al., 2011; Chapon et al., 2012). A small localized study found more bacterial OTUs in the high-radiation samples than in the low-radiation samples (Theodorakopoulos et al., 2017). Other studies, however, have reported a lower diversity of bacteria from the most highly radio-contaminated sites. For example, the diversity of cultured bacteria sampled in the 10-km zone around the Chernobyl Nuclear Power Plant was two orders of magnitude lower than in control, non-contaminated areas (Romanovskaia et al., 1998). Similarly, the diversity of soil bacterial communities was lower in samples from the most highly radio-contaminated areas within Fukushima (Ihara et al., 2021). There are some possible factors behind this large variation in microbial diversity results across studies. The combination of using a metabarcoding technique with high-throughput sequence data (Illumina NovaSeq) and extensive environmental sampling allowed us to conduct a comprehensive evaluation of the bacterial communities in the area. This methodological implementation led to a massive improvement in the detection of bacteria compared to previous studies, including lineages that are rare or hard to culture (Woo et al., 2008; Joos et al., 2020). In addition, the bacterial communities in the area are likely to differ between wetlands and terrestrial environments which have been targeted in previous studies (e.g. Romanovskaia et al., 1998; Chapon et al., 2012; Theodorakopoulos et al., 2017). Furthermore, over thirty years have passed since the accident in Chernobyl, and the chronic exposure to low-dose radiation may have facilitated the proliferation of bacteria resistant to radiation (already suggested by Zavilgelsky et al., 1998), which may explain why some of the studies conducted closer in time to the nuclear accidents in Chernobyl (Romanovskaia et al., 1998) or Fukushima (Ihara et al., 2021) found lower bacterial diversity measures. Bacterial adaptation to radioresistance can result from relatively small genetic changes affecting DNA repair and metabolic functions (DeVeaux et al., 2007; Harris et al., 2009; Byrne et al., 2014), and these evolutionary responses are much more likely to appear with increasing time after radiation exposure.

The composition of bacterial communities in Chernobyl wetlands was affected by diverse environmental factors, including radiation. Overall, soil communities were characterized by the abundance of Actinobacteriota, a phylum typically dominant in this environment (Hill et al., 2011), abundant also in previous studies in Chernobyl (Theodorakopoulos et al., 2017). Water communities were dominated by Proteobacteria and Bacteroidota, two of the most common phyla of bacteria in freshwater environments (Newton et al., 2011). When examined using Bray-Curtis dissimilarity, the composition of bacterial communities in water was affected by the amount of dissolved organic matter and water temperature, both of which have been previously recognized as potential drivers of bacterial community structure (e.g. Judd et al., 2006; Hall et al., 2008). Soil and sediment community structure was mostly influenced by radiation levels. Although pH has been shown to be a key determinant of the composition of bacteria communities in Chernobyl (Antwis et al., 2021), and elsewhere (e.g. Griffiths et al., 2011; Zhalnina et al., 2015), we did not detect any significant effect of pH in any of the sampled environments. When using UniFrac phylogenetic distances, the effect of radiation was strongly significant in soil and water, indicating that radiation has a strong effect on the phylogenetic composition of the microbiome in these environments.

The main influence of radiation on the bacterial communities of Chernobyl was detected when examining the taxonomic composition of the different microbiomes. In particular, we identified several bacterial families with higher abundances in sites with higher radioactivity. Among them, many taxa are reported as common in radioactive environments, with some being able to reduce uranium and other radioactive metals. For example, several of the taxa that were more abundant in water with high radiation levels included *Rhodocyclaceae*, *Smithellaceae*, *Geobacteraceae*, Synthrophales and *Desulfobaccaceae*. *Rhodocyclaceae* is a family that includes UV-radiation resistant members (Han et al., 2020) and has been detected also in the Handford 300 area, a former complex for radioactive fuel manufacture (Converse et al., 2015). *Geobacteraceae* are metal-reducers that have been frequently found in areas rich in uranium (Suzuki et al., 2005; Simonoff et al., 2007; N’Guessan et al., 2010; Zachara et al., 2013, Sutcliffe et al., 2018). *Smithellaceae*, *Desulfobaccaceae* and Synthrophales have also been detected in disposal sites for liquid radioactive waste and experimental areas with radionuclide contamination (Nazina et al., 2010, Vikmna et al., 2019, Gihring et al., 2021). In our study, among the microbes that were more abundant in high radiation sediment samples were *Anaerolineae*, *Microbacterium* and the archaea Thermoplasmata. These three taxa have also been previously associated with uranium-rich soils (Mondani et al., 2011), groundwater from nuclear waste depositories (Nedelkova et al., 2007), and radioactive legacy sites (Vazquez-Campos et al., 2021), respectively. *Prolixibacteraceae,* which were associated with high soil radiation levels in our study, have also been detected in bogs with high levels of radioactive selenium, cesium, thorium and uranium (Lusa & Bomberg, 2021). Several of the groups that were abundant in high radiation soils, e.g. *Lentimicrobiaceae*, are slow-growing bacteria associated with polluted environments. Radioresistant bacteria often have slow growth cycles, diverting their resources from growth to DNA repair and stress resistance mechanisms (Zakrzewska et al., 2011). For some of the other taxa, we still have little information about their ecological requirements, in particular any potential resistance to radiation. Further studies on the effects of radioactivity on specific microbial taxa are therefore necessary.

In summary, we detected rich and diverse microbial communities in the wetlands of the Chernobyl Exclusion Zone, regardless of radiation levels. Multiple bacterial and archaeal taxa had higher abundance in the high-radiation sites, many of which were previously reported to thrive in other areas with high radioactivity. By using extensive field sampling together with metabarcoding and high-throughput sequencing, our study significantly contributes to our understanding of the drivers that affect the diversity and composition of bacterial communities in radio-contaminated environments. The host-associated microbiomes of multicellular organisms (plants and animals) that live in these highly radioactive areas may be directly sourced from the microbial communities present in the local environment (as suggested by Jones et al., 2004). Thus, an improved knowledge of how environmental microbiomes respond to a gradient of radioactivity is therefore crucial to forecast the future functionality and re-naturalization potential of radiocontaminated environments.

## ACKNOWLEDGEMENTS

We are thankful to Sergey Gaschack for his invaluable help with field sampling and radiation evaluation, and to the staff of the Chornobyl Center for Nuclear Safety, Radioactive Waste and Radioecology (Slavutych, Ukraine) for their support during field research. We thank AllGenetics & Biology SL for help with DNA metabarcoding work and great analytical support. Katerina Guschanski provided valuable comments on initial stages of the study. This work was supported by projects from the Helge Ax:son Johnsons Stiftelse to PB, and Swedish Radiation Protection Agency-SSM (SSM2018-2038) and Carl Tryggers Foundation (CT 16:344) to GO. EV was supported by a fellowship from the Swedish Research Council (2020-00259), PB by a Carl Tryggers Foundation scholarship (CT 16:344) and by a Marie Sklodowska-Curie fellowship (METAGE-797879), and GO by the Spanish Ministry of Science, Innovation and Universities “Ramón y Cajal” grant RYC-2016-20656.

## AUTHOR CONTRIBUTIONS

G.O. conceived the study. P.B. and G.O. conducted field surveys and sample collection. E.V. analyzed the data. E.V. and G.O. wrote the manuscript with significant contributions from P.B.

## DATA AVAILABILITY STATEMENT

Detailed information of all sampling sites is available in the supplementary information. The sequence reads have been uploaded to EMBL-EBI ENA and will be made public upon acceptance of the manuscript.

---

**Table S1.**
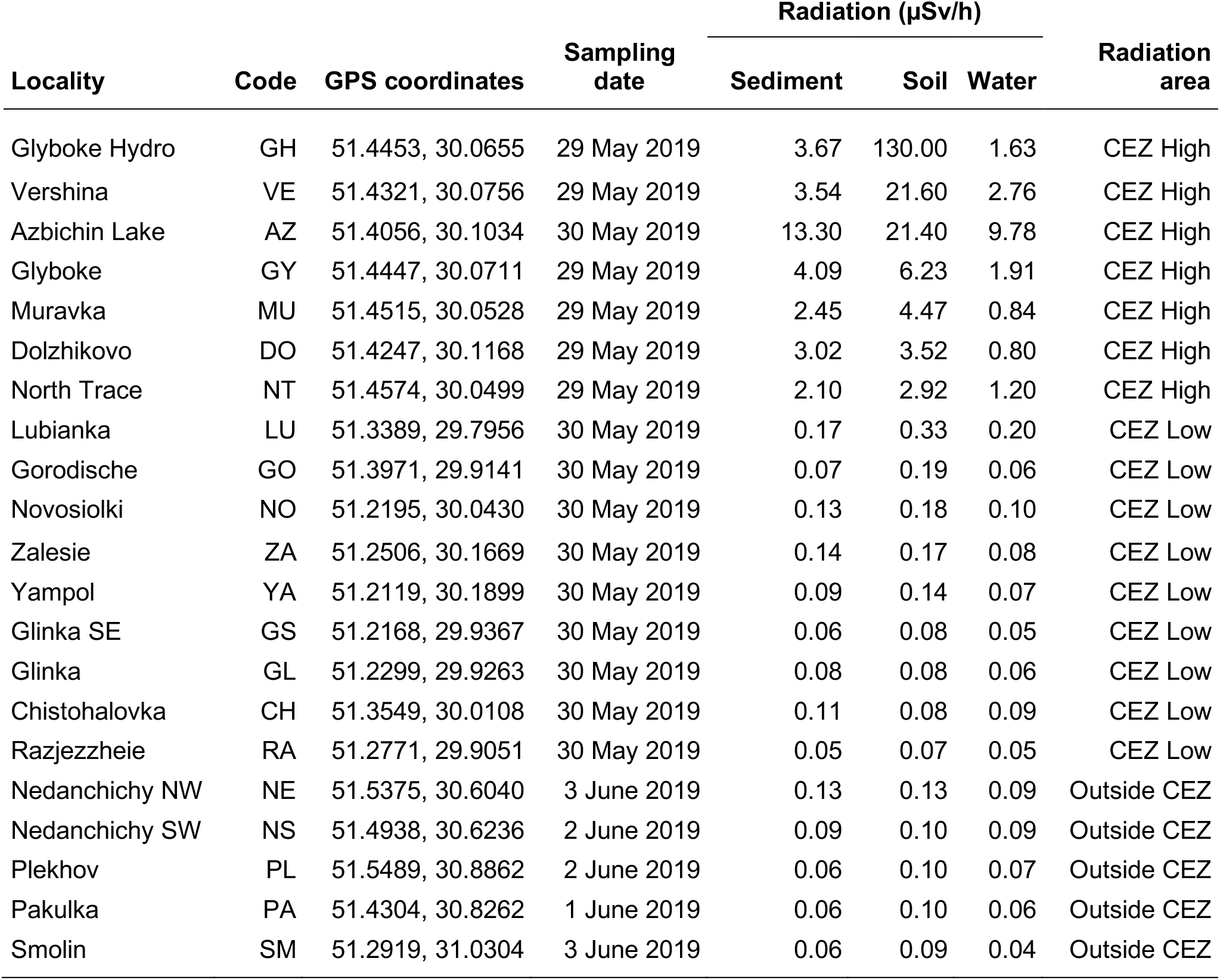
Geographic coordinates (latitude and longitude), sampling date, current levels of radiation (in water, pond sediment and shore soil), and radiation area of the wetlands included in the study. CEZ: Chernobyl Exclusion Zone.

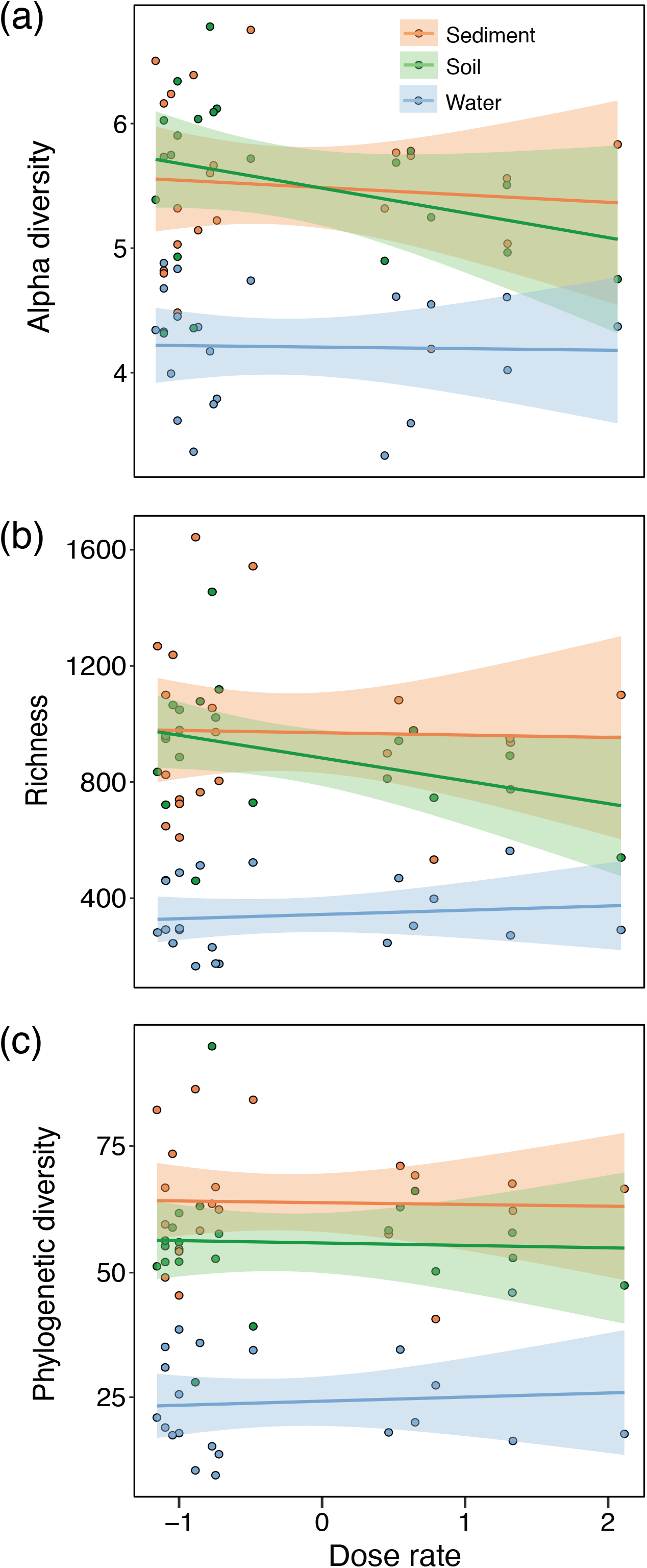

